# Lipid scrambling via TMEM16F mediates the formation and release of apoptotic vesicles

**DOI:** 10.1101/2025.05.14.654002

**Authors:** Trieu Le, Yu Meng Li, Amra Saric, Joost C.M. Holthuis, Sergio Grinstein, Spencer A. Freeman

## Abstract

The ubiquitous and highly conserved programmed cell death pathways that are essential for tissue development and homeostasis are accompanied by distinct morphological alterations. Apoptotic cells undergo fragmentation that is concomitant with the exposure of phosphatidylserine (PS) on the membrane surface. Large fragments, called apoptotic bodies, as well as much smaller and more numerous vesicles are released. While the molecular mechanisms underlying apoptotic body formation have been explored, much less is known about vesicle biogenesis. We used an inducible, active form of TMEM16F to determine the role of lipid scrambling in vesiculation, separately from other apoptotic signaling events. Plasmalemmal lipid scrambling sufficed to release apoptotic-like vesicles without causing changes in cytosolic calcium or the submembrane cytoskeleton. The scrambled bilayer showed pronounced segregation of exofacial lipids and redistribution of apparent cholesterol to the inner leaflet. The clustering of raft-associated components with bulky headgroups –typified by glycophosphatidylinositol-linked proteins– formed domains of outward (convex) curvature, while regions of accumulation of phosphatidylethanolamine (PE) generated inward (concave) curvature that facilitated the scission of vesicles. Thus, scrambling of plasma membrane lipids suffices to induce regions of acute membrane curvature and facilitates detachment of vesicles analogous to those released from the surface of apoptotic cells.

## Introduction

Apoptosis remains the best characterized form of programmed cell death, playing essential roles in tissue renewal, development, and immunity[1]. Two hallmarks of apoptotic cells, the exposure of phosphatidylserine (PS) on their outer surface and their fragmentation[2], foster the detection and clearance of the dying cells[3]. Apoptotic cells release fragments of widely varying size: apoptotic bodies (ApoBDs) ranging from 1-5 µm in diameter and the much smaller apoptotic vesicles (ApoVs) of 0.1-1 µm in diameter[4]. ApoBDs and ApoVs differ not only in their size but also in their composition and therefore play distinct roles in signaling and clearance of the dying cells. In addition to PS, the smaller ApoVs are enriched in certain surface markers including tetraspanins, C1q, and CD44, while excluding other transmembrane proteins like CD45[5]. By contrast, ApoBDs contain many of the components of the membrane and cytosol[6–8].

The membrane blebbing and fragmentation that accompany apoptosis involve remodelling of the actin cytoskeleton: cleavage of some structural components of the membrane skeleton occurs in conjunction with myosin-supported contractility. These changes are associated with drastic changes in cytosolic calcium concentration. Calcium-activated proteases and phosphatases have been implicated in the stimulation of Rho-associated protein kinase (ROCK), p21-activated kinase (PAK), and gelsolin, that regulate cytoskeletal contraction[9]. Calcium-activated proteases, notably calpain and caspases, also promote the cleavage of spectrin[10] and kinases that normally activate ERM proteins[11], structural components that connect the actin cortex to the plasma membrane. Spectrins and ERM proteins are also controlled by anionic phospholipids whose abundance and transmembrane distribution are stringently regulated by calcium: the cation can stimulate phospholipases and activates lipid scramblases directly (e.g. TMEM16F)[12] or, in the case of XKR8, through stimulation of caspase-3[13].

Despite obvious differences in their size and composition, it remains unclear whether ApoBDs and ApoVs arise from the same mechanistic process. It is generally accepted that ApoBD formation involves the cytoskeletal remodelling and contraction detailed above, but whether these are required for ApoVs detachment is not evident. Clearly, the contribution of individual components to the two modes of fragmentation needs to be analyzed separately. In this context, it was recently reported that the deletion of a single lipid scramblase, TMEM16F, suppresses the membrane blebbing induced by elevated cytosolic calcium[14, 15]. It was therefore of interest to establish whether TMEM16F is necessary or even sufficient to generate ApoVs. Isolating the contribution of this scramblase is challenging, due to the pleiotropic effects of large calcium changes such as those that accompany apoptosis. However, recent molecular characterizations of TMEM16F provide a potential means of overcoming these limitations: targeted amino acid substitutions within its hydrophobic pore have enabled the generation of scramblase variants that are active at physiological (resting) calcium concentration[16]. These advances provided us with an opportunity to systematically analyze the role of transmembrane lipid distribution in the formation of ApoV and ApoBD, separately from the other processes that underlie apoptosis.

## Results

### Activation of TMEM16F is sufficient to drive vesiculation

We expressed constitutively active forms of TMEM16F to assess the consequences of plasmalemmal lipid scrambling without elevating cytosolic [Ca^2+^]. These were generated by mutating a critical tyrosine residue (Y563) found in the lining of the pore, which occludes lipid transport in the wildtype scramblase when cytosolic [Ca^2+^] is in the low nM range, i.e. in unstimulated healthy cells (Fig 1A)[16]. Like the wildtype (WT) scramblase, these mutant forms of TMEM16F (Y563A or Y563Q, tagged with mCherry) were delivered to the cell surface when transiently expressed in primary human fibroblasts (Fig 1B). However, unlike the WT TMEM16F, the Y563A and Y563Q versions induced lipid scrambling when cytosolic [Ca^2+^] was maintained at resting levels, as indicated by the appearance of PS on the outer leaflet of the membrane, detected using Alexa647-conjugated Annexin V (AnV) (Fig 1B). Remarkably, staining of PS also revealed extensive vesiculation of the membrane. These vesicles were also clearly visible using the solvochromic dye FM4-64, indicating that they were not induced by AnV; moreover, unstained vesicles were also detectable electronically (see below). The vesicles had two important features. First, most appeared to be ≤1 μm in diameter, consistent with published measurements for ApoVs. Second, the vesicles stained more brightly with AnV than the bulk of the membrane of the transfected cells (Fig 1B).

**Figure 1.**
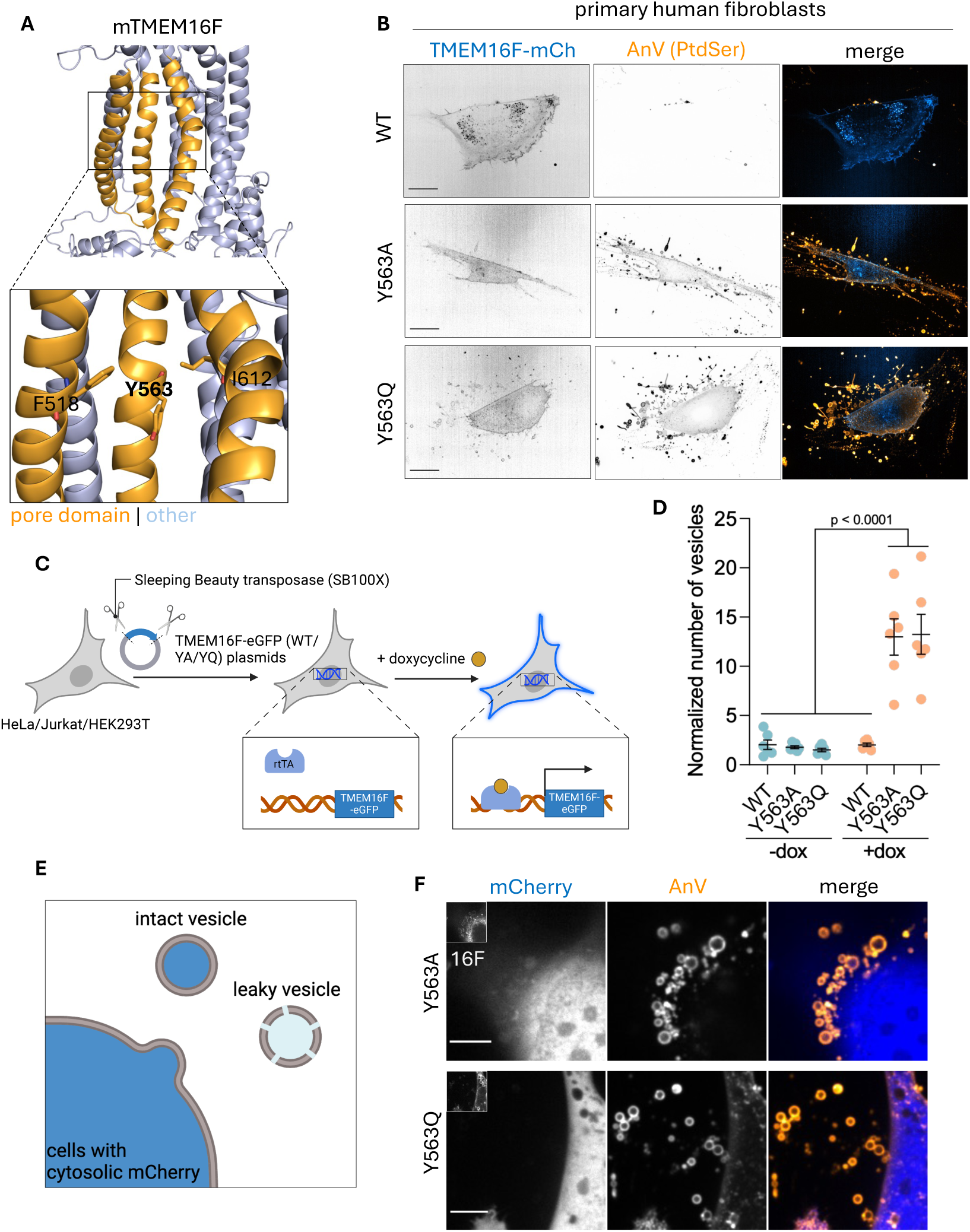
Lipid scrambling by TMEM16F drives vesiculation. A) Homology model of the open (Ca^2+^-activated) state of mTMEM16F, derived using the Ca^2+^-bound structure of nhTMEM16 (PDB: 4WIS). B) Expression of the TMEM16F constitutively active mutants, Y563A and Y563Q, induces the formation of vesicles. WT, Y563A and Y563Q mCherry (mCh) transiently expressed in primary human fibroblasts. A single plane of the TMEM16F-mCh signal and a z stack of the Annexin V (AnV) signal are shown. Scale bar: 20 μm. C) Diagram of inducible cell system. D) Quantification of vesicles produced by HeLa cells with or without doxycycline (dox). E) Model of vesicle generation. F) TMEM16F-induced vesicles do not retain cytosolic mCherry. Scale bar: 5 μm.

To perform quantitative determinations of lipid scrambling and vesiculation across cell populations, we generated cells stably expressing the active forms of TMEM16F. This was achieved by integrating doxycycline-inducible WT, Y563A and Y563Q TMEM16F tagged with eGFP into HeLa, Jurkat, and HEK293T cells using the Sleeping Beauty transposase system (Fig 1C). Acute inducibility by doxycycline was required because cells constitutively expressing the mutant scramblases were not viable for long-term culture. Once induced, the WT and mutant scramblases trafficked similarly to the plasma membrane. However, as was the case for the transient transfectants, overnight induction caused lipid scrambling and vesiculation in the case of the mutants, but not the WT TMEM16F. The occurrence of vesiculation could be quantified by electronic sizing using a Coulter counter set to discern particles of diameter between 1-5 μm. As shown in Figure 1D, HeLa cells induced to express active TMEM16F (Y563A- or Y563Q-eGFP) released such particles (vesicles) into the medium in abundance when compared to their non-induced counterparts or to cells expressing the WT form of the scramblase.

### Scrambling-induced vesicles are leaky

The previous observations indicated that vesicles produced upon activation of TMEM16F stained brightly with AnV. This could indicate that PS is enriched on the outer leaflet of the vesicles compared to the bulk plasma membrane or, alternatively, that the vesicles are sufficiently permeable to AnV to enable the probe to access PS on both sides of the bilayer. To assess whether the vesicles become leaky during their biogenesis, we co-expressed a cytosolic indicator, mCherry, together with the active TMEM16F mutants fused to GFP. As illustrated in Figure 1E, most of the AnV-positive vesicles generated by the active scramblases failed to retain mCherry in their lumen, implying that they are leaky to relatively large molecules (≈30 kDa), likely including AnV. That the vesicles are leaky was confirmed using FM4-64, which in membrane impermeant. This amphiphilic dye, which fluoresces brightly when inserted into lipid bilayers, stained most of the Y563A- or Y563Q-induced vesicles more brightly than the bulk membrane of the transfected cell (S Fig 1), suggesting that FM4-64 partitioned into both aspects of their bilayer.

### Lipid scrambling-induced vesiculation is largely independent of extracellular Ca^2+^

In addition to serving as a Ca^2+^-activated lipid scramblase, TMEM16F has also been shown to function as a small conductance non-selective cation channel[17]. Because the cations traverse the same pore used to scramble lipids, it was conceivable that the constitutively active Y563A or Y563Q mutants would promote extracellular Ca^2+^ influx; this might result in a cytosolic [Ca^2+^] increase that would confound the interpretation of the vesiculation process. To assess this possibility, we estimated the cytosolic [Ca^2+^] using GCaMP6s. To make them comparable regardless of the expression level of the sensor, the resting signals –measured 16 h after inducing the expression of the mutants– were normalized to the maximal fluorescence. The latter was recorded after addition of saturating concentrations of ionomycin, a calcium ionophore that was added at the end of each determination. As illustrated in Figure 2A, the resting [Ca^2+^] was similar in cells expressing WT and the Y563A or Y563Q forms of the scramblase. These findings imply that an elevation of cytosolic [Ca^2+^] was not involved in the generation of the vesicles induced by the mutant scramblases. This conclusion was supported by experiments where induction of the scramblases was carried out in media devoid of Ca^2+^. Vesicle release under the various conditions used was quantified by electronic sizing, as described above. We noted a modest decrease in the production of vesicles upon Ca^2+^ removal; however, the active TMEM16F cells still showed pronounced vesiculation even in the absence of extracellular Ca^2+^ (Fig 2B).

**Figure 2.**
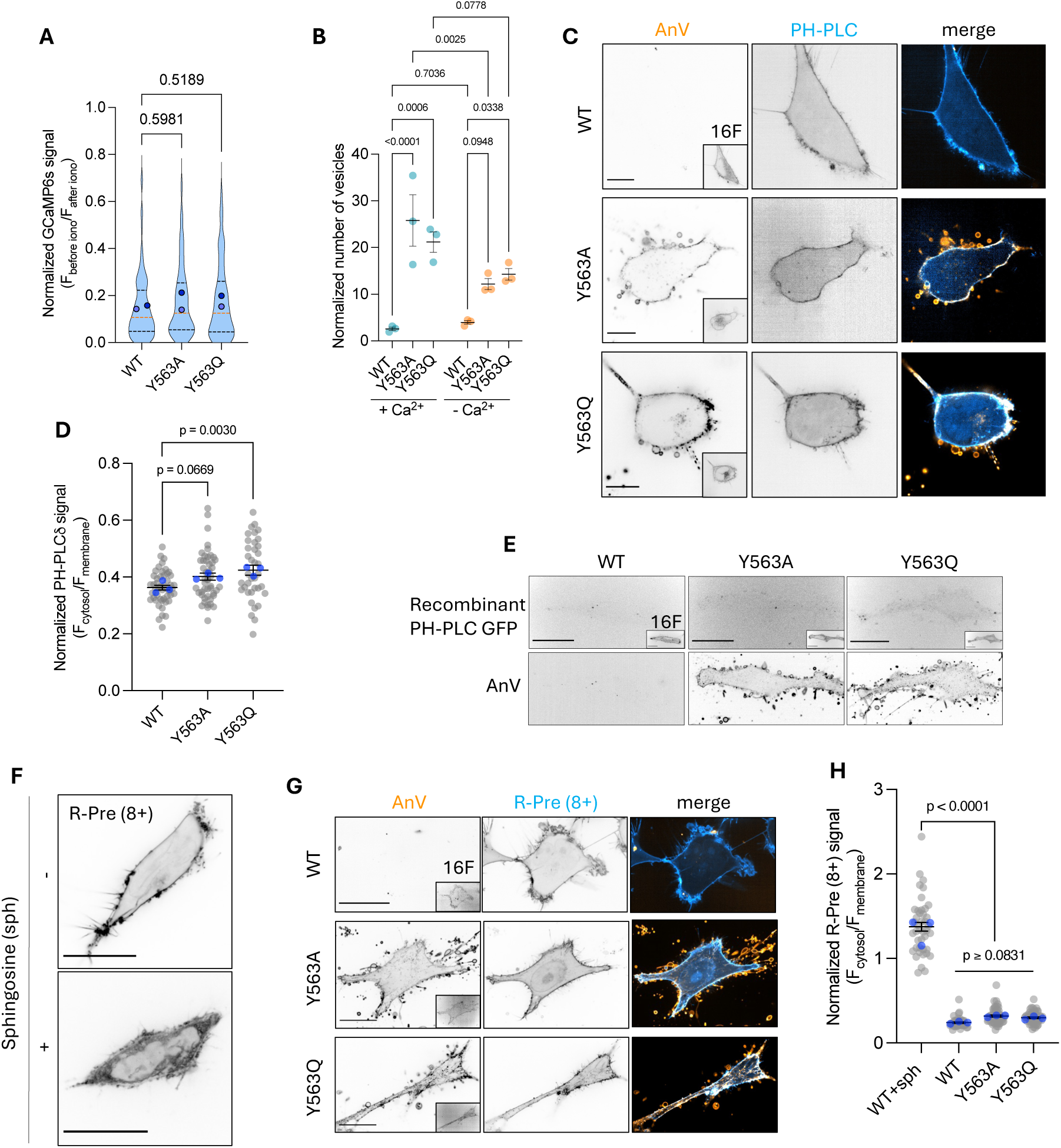
TMEM16F scrambling activity does not change phosphoinositides or membrane charge of the inner leaflet. A) [Ca^2+^] determined with GCaMP6s in the cytosol of primary human fibroblasts expressing mCherry-tagged WT, YA or YQ TMEM16F. Signals were normalized to the maximum, i.e. after the addition of ionomycin. B) Quantification of vesicle number in doxycycline-induced stable cells in the presence or absence of external Ca^2+^. C) WT and YQ TMEM16F expressed with PH-PLC-mCherry in HeLa cells with exofacial PS labeled with AnV. D) Quantification of F_cytosol_/F_membrane_ for PH-PLC-GFP signal in *C*. E) WT, YA or YQ TMEM16F-expressing HeLa cells labeled with recombinant PH-PLC-GFP. F) A representative R-Pre 8+-expressing cell before and after addition of sphingosine. G) WT, YA and YQ TMEM16F co-expressed with R-Pre 8+ in HeLa cells. H) Quantification of F_cytosol_/F_membrane_ R-Pre 8+ from *F* and *G*. All scale bars: 20 μm.

### Activation of TMEM16F does not change PtdIns(4,5)P_2_ levels or the negative surface charge of the plasmalemmal inner leaflet

We next investigated whether weakening of the interaction between the membrane and the associated cytoskeleton was responsible for scramblase-induced vesiculation. In this regard, when the integrity of the membrane-associated skeleton is compromised, the outward hydrostatic pressure exerted by the existing slight osmotic imbalance was reported to cause membrane blebbing[18]. Phosphoinositides, especially PtdIns(4,5)P_2_, play a key role in fastening the cytoskeleton to the membrane. Therefore, depleting PtdIns(4,5)P_2_ from the inner leaflet of the plasmalemma through scrambling could conceivably weaken the membrane-cytoskeleton interaction. For this reason, we investigated whether the active TMEM16F mutants influenced the transmembrane distribution of PtdIns(4,5)P_2_. This was achieved by expressing the pleckstrin-homology domain of phospholipase C (PH-PLC), a PtdIns(4,5)P_2_ biosensor. As shown in Figure 2C and quantified in Figure 2D, scrambling of the plasma membrane by the active TMEM16F mutants did not visibly alter the abundance of the lipid on the cytosolic-facing leaflet. To complement these observations, we also tested whether PtdIns(4,5)P_2_ became exposed on the outer leaflet. To this end, we added recombinant PH-PLC to the extracellular medium bathing WT- or mutant-expressing cells. The probe failed to detect PtdIns(4,5)P_2_ in all instances (Fig 2E), a finding compatible with the retention of the lipid in the inner leaflet.

The inner leaflet of the plasmalemma is the most negatively charged cytosol-facing membrane, due to its elevated content of anionic lipids, mainly PS, phosphatidic acid, and phosphoinositides[19]. Lipid scrambling stands to impact the surface charge of the inner leaflet, which could play a role in vesiculation. We used a probe for membrane surface charge to assess this possibility. The probe used, R-Pre (8+), binds preferentially to highly negative membranes by virtue of its prenylated and polycationic tail[19]. As proof of principle, we demonstrated that R-Pre (8+), which largely partitions to the plasma membrane when expressed in otherwise untreated cells, dissociates immediately upon addition of sphingosine, a positively-charged 18-carbon amino alcohol that readily incorporates into the bilayer and masks its negative surface charge (Fig. 2F). In contrast, and as shown in Figure 2G and quantified in Figure 2H, the preferential recruitment of R-Pre (8+) to the plasma membrane persisted in lipid-scrambled cells.

The release of a modest fraction of the probe in the case of the mutants is consistent with the concomitant PS scrambling, but the persistent negativity likely reflects the retention of the multivalent PtdIns(4,5)P_2_, which has a much greater negative charge (approx. -4 at physiological pH) and charge density than PS or phosphatidic acid (charge = -1). It is noteworthy that neither the R-Pre (8+) nor the PH-PLC probes expressed in the mutants were detected in the vesicles (Fig 2C, 2G). While able to bind to membranes, both of these probes dissociate rapidly into the cytosol. Their absence from the vesicles is consistent with the conclusion that the latter are leaky. Together, the preceding results suggest that neither loss of phosphoinositides nor depolarization of the negative surface charge of the inner leaflet play a significant role in vesiculation.

### Activation of TMEM16F does not visibly alter the membrane cytoskeleton

As described above, formation of ApoBDs requires the appearance of discontinuities in the connection of the plasma membrane with the cytoskeleton, and contraction of the latter: cleavage of cytoskeletal components and activation of myosins underlie these effects, respectively. The role of the cytoskeleton and its associated myosins in the production of ApoVs is less clear. We therefore investigated whether the cytoskeleton was involved in the vesiculation triggered by TMEM16F. We first used LifeAct, an F-actin binding probe, in cells that also expressed WT or the active mutants of TMEM16F. As shown in Figure 3A, we found that the fluorescence intensity of cortical LifeAct, a coarse measure of F-actin association with the plasma membrane, was similar in WT and mutant cells. Quantitation of optical midsections of multiple cells (Fig. 3B) confirmed that the cortical F-actin remained unaltered in vesiculating cells.

**Figure 3.**
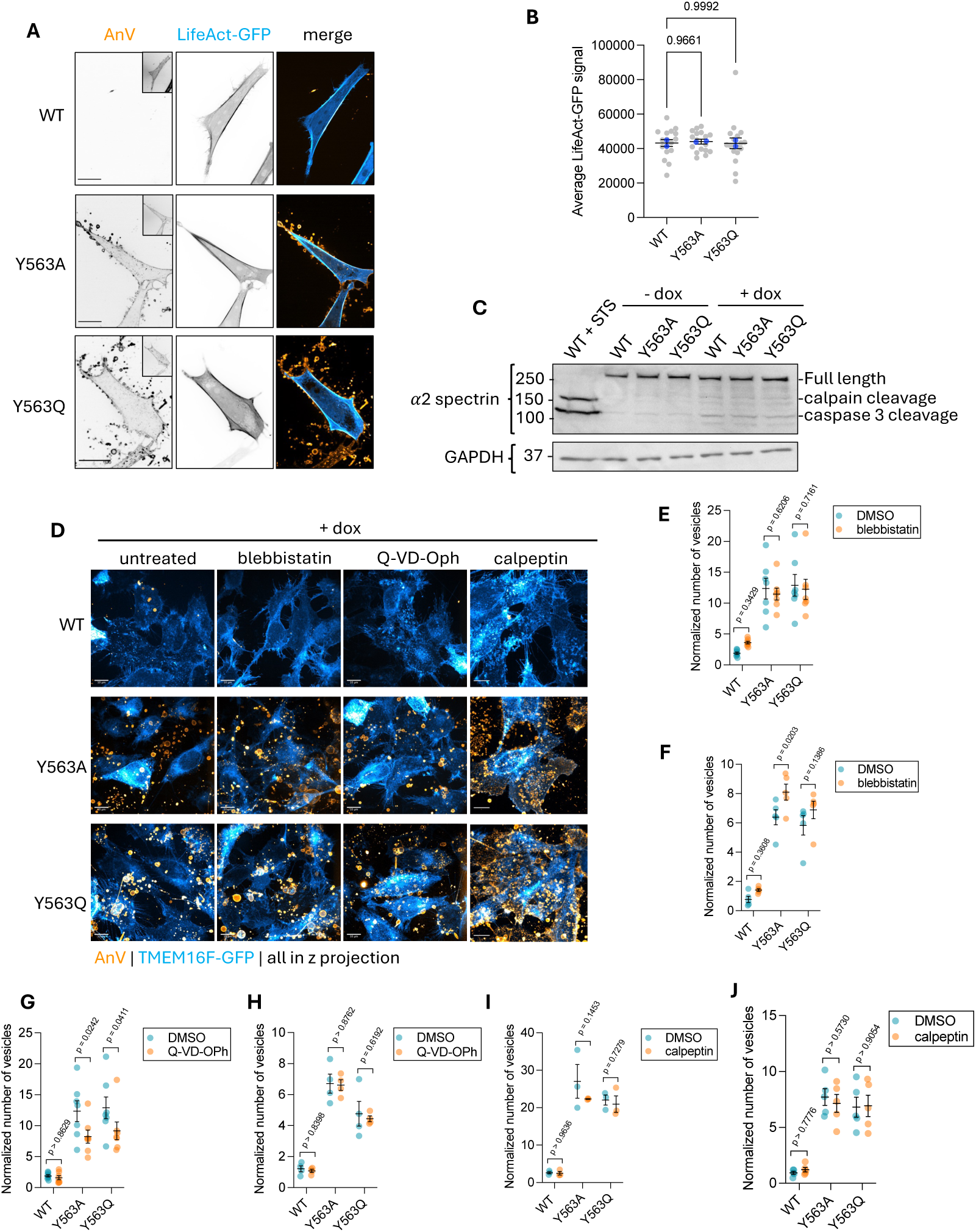
Lipid scrambling-induced vesiculation is independent of cytoskeletal remodeling. A) WT/YA/YQ TMEM16F-mCherry and LifeAct-GFP-expressing HeLa cells were labelled live for exposed PS AnV. B) Analysis/quantification of F-actin in induced cells. C) Western blotting for α2 spectrin in HeLa cells. Intact spectrin and products of its degradation indicated. D) WT/YA/YQ inducible HeLa cells treated with blebbistatin, a caspase inhibitor (Q-VD-OPh) or calpeptin labelled with AnV. E) Vesicle quantification upon blebbistatin treatment of inducible HeLa cells. F) Vesicle quantification in inducible Jurkat cells. G) Vesicle quantification upon caspase inhibitor treatment of inducible HeLa cells. H) Vesicle quantification upon caspase inhibitor treatment of inducible Jurkat cells. I) Vesicle quantification in calpeptin treated inducible HeLa cells. J) Vesicle quantification in calpeptin treated of inducible Jurkat cells. All scale bars: 20 μm

We next determined if α2 spectrin, a critical component of the cytoskeleton that is cleaved by calpains and caspases during apoptosis, was affected upon bilayer scrambling. To validate the sensitivity of the immunoblotting procedure used to detect spectrin breakdown, *bona fide* apoptosis was induced in WT cells with staurosporine. As illustrated in Figure 3C, two major cleavage products were readily observed in these cells, while the full-length α2 spectrin virtually disappeared. In sharp contrast, the intact α2 spectrin persisted in cells where marked vesiculation was induced by expression of the active scramblases.

These findings suggest that cytoskeletal cleavage by caspases and/or calpain is not involved in the generation of the vesicles. This concept was tested further by incubating the cells with potent inhibitors of caspases or calpain. Neither Q-VD-OPh (a pan-caspase inhibitor), nor calpeptin (a calpain inhibitor) prevented vesicle formation by the active scramblases (Figs. 3D, G-J). Further, we did not observe differences in the vesiculation of the cells when myosin II was inhibited with blebbistatin (Fig. 3D-F), which is well documented to prevent ApoBD formation[20]. Jointly, these data indicate that cytoskeletal disruption and contractility are not responsible for vesiculation during bilayer scrambling, further reinforcing the finding that elevations in [Ca²⁺] are not required for this process.

### Sphingomyelin translocation and hydrolysis are not involved in scrambling-induced vesiculation

In endolysosomes, membrane damage is associated with lipid scrambling. The resulting exposure of luminal sphingomyelin to cytosolic sphingomyelinases facilitates resealing of the damaged organelles[21]. Conceivably, scrambling of plasmalemmal sphingomyelin with its consequent exposure to cytosolic neutral sphingomyelinases might contribute to the vesiculation induced by the Y563A or Y563Q variants of TMEM16F. Indeed, cleavage of the relatively large phosphocholine headgroup of sphingomyelin to produce ceramide –a conical lipid that accommodates/generates inward (negative, concave) curvature– is thought to enable vesiculation [21, 22]. To test the possible involvement of sphingomyelin in vesiculation we used a biosensor derived from equinatoxin, EqtSM-GFP. When expressed in HeLa cells, EqtSM-GFP was capable of sensing sphingomyelin exposure to the cytosol when lysosomes were ruptured with LLOMe (Fig. 4A). That the biosensor specifically detected sphingomyelin in such experiments was confirmed using HeLa cells edited to delete expression of both sphingomyelin synthases, SMS1 and SMS2: no redistribution of EqtSM-GFP was observed in these cells when lysosomes were ruptured (Fig. 4B). We then expressed EqtSM-GFP together with the TMEM16F WT and active mutants to compare the distribution of sphingomyelin with or without lipid scrambling. As shown in Figure 4C, we did not detect any enrichment of sphingomyelin in the inner leaflet of the plasma membrane following TMEM16F activation.

**Figure 4.**
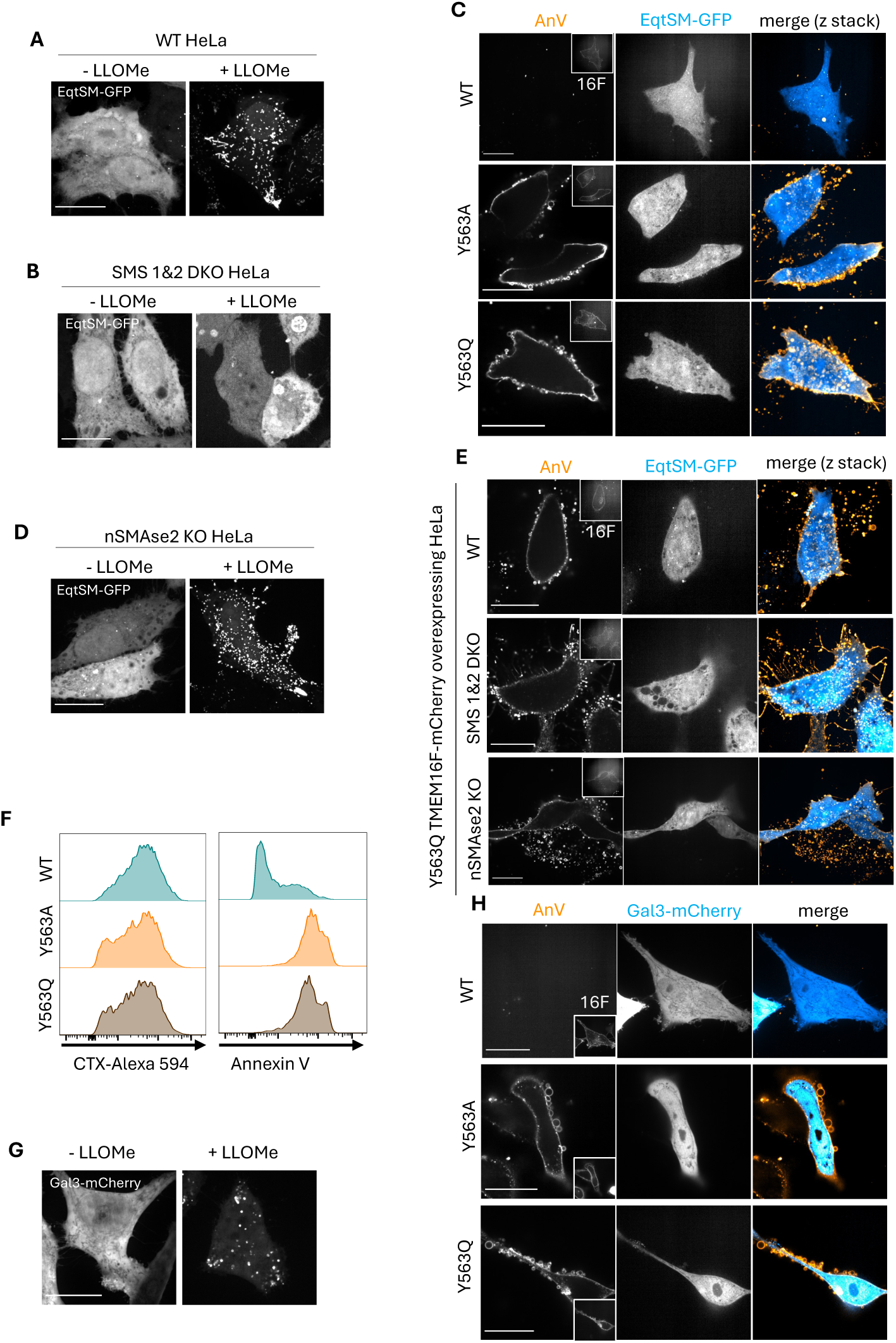
TMEM16F-induced vesiculation is independent of sphingomyelin. A) WT HeLa cells expressing EqtSM-GFP treated with LLOMe. B) SMS 1/2 DKO HeLa cells expressing EqtSM-GFP, treated with LLOMe. C) WT/YA/YQ TMEM16F-mCherry expressed together with EqtSM-GFP in WT-HeLa cells, labeled AnV. D) nSMAse2 KO HeLa expressing EqtSM-GFP, treated with LLOMe. E) Y563Q TMEM16F-mCherry expressed with EqtSM-GFP in WT, nSMAse2 KO, SMS1/2 DKO or nSMAse2 KO HeLa cells, labeled with AnV. F) Flow analysis of WT/YA/YQ TMEM16F inducible Jurkat cells labeled with AnV and cholera toxin B. G) WT HeLa cells expressing Gal3-mCherry treated with LLOMe. H) Galectin-3 mCherry expressed in WT/YA/YQ TMEM16F inducible HeLa cells, labeled with AnV. All scale bars: 20 μm

Failure to detect EqtSM-GFP binding to the membrane may have been due to rapid conversion of scrambled sphingomyelin to ceramide by the cytosolic neutral sphingomyelinase (nSMAse2). Cells devoid of nSMAse2 were used to evaluate this possibility. As shown in Figure 4E, sphingomyelin was not detected on the inner aspect of the plasma membrane of nSMAse2-deficient cells expressing the active scramblase, despite the fact that the EqtSM-GFP probe is still effective in these cells, as shown in Fig. 4D. These findings indicate that sphingomyelin is not scrambled by TMEM16F and hence is not required for vesiculation. This was tested directly by expressing the Y563Q mutant in cells lacking both SMS1 and SMS2. These double-knockout cells underwent seemingly normal vesiculation despite lacking sphingomyelin (Fig. 4E). Vesiculation also occurred in nSMAse2-deficient cells (Fig. 4E), confirming that generation of ceramide through sphingomyelin hydrolysis is not involved in vesicle formation.

### Glycolipids are not scrambled by TMEM16F

In addition to the phospholipids transported through lipid scramblases, lipids with large headgroups (e.g. glycolipids) have also been reported to be scrambled across bilayers by the TMEM16 family of scramblases, at least in liposomes[23]. A prominent glycolipid found in the outer leaflet of cells is the ganglioside, GM1. Using the B-subunit of cholera toxin as a probe, we found only a minute decrease in the abundance of the ganglioside in a fraction of the cells, which may have resulted from loss via vesiculation or exofacial hydrolysis (Fig. 4F). If occurring at all, scrambling of GM1 would be minimal. Other glycolipids could nevertheless be involved. We therefore expressed galectin-3 (Gal-3) fused to mCherry in the HeLa cells in order to determine if any β-galactoside-containing glycolipids were exposed to the cytosol. Of note, Gal-3-mCherry becomes enriched in lysosomes upon their rupture with LLOMe (Fig. 4G), validating its ability to detect β-galactoside moieties in our cells. As shown in Figure 4H, however, we failed to detect any appearance of Gal-3-binding lipids on the plasma membrane of cells expressing the active forms of TMEM16F. Together, these data suggest that scrambling of glycolipids, like sphingolipids, does not contribute to vesiculation.

### The redistribution of GPI-anchored proteins, cholesterol and PE alters the membrane topology in a manner that supports vesiculation

How does scrambling of the plasma membrane bilayer ultimately result in its vesiculation? Because neither cytoskeletal contractility nor cytosolic enzymes seemed to be implicated, we hypothesized that intrinsic features of the scrambled bilayer may generate curvature that favors vesicle formation. This was initially suggested by the observation that GPI-anchored proteins, visualized by expressing glycophosphatidylinositol-linked GFP (GPI-GFP), showed clear enrichment in the forming vesicles (Fig. 5A). GPI-anchored proteins normally reside together with cholesterol in raft-like regions of the outer leaflet[24]. This prompted us to analyze the distribution of cholesterol in cells that had undergone scrambling. The distribution of exofacial cholesterol was visualized using recombinant D4-GFP[25]. This probe binds primarily free cholesterol but not cholesterol complexed to sphingomyelin[26]. D4-GFP revealed that cholesterol was largely depleted from the main body of the scrambled cells and instead was enriched on the vesicles (Fig. 5B). These observations are consistent with the notion that cholesterol redistributes laterally on the outer leaflet of the membrane, replicating the distribution of GPI-GFP.

**Figure 5.**
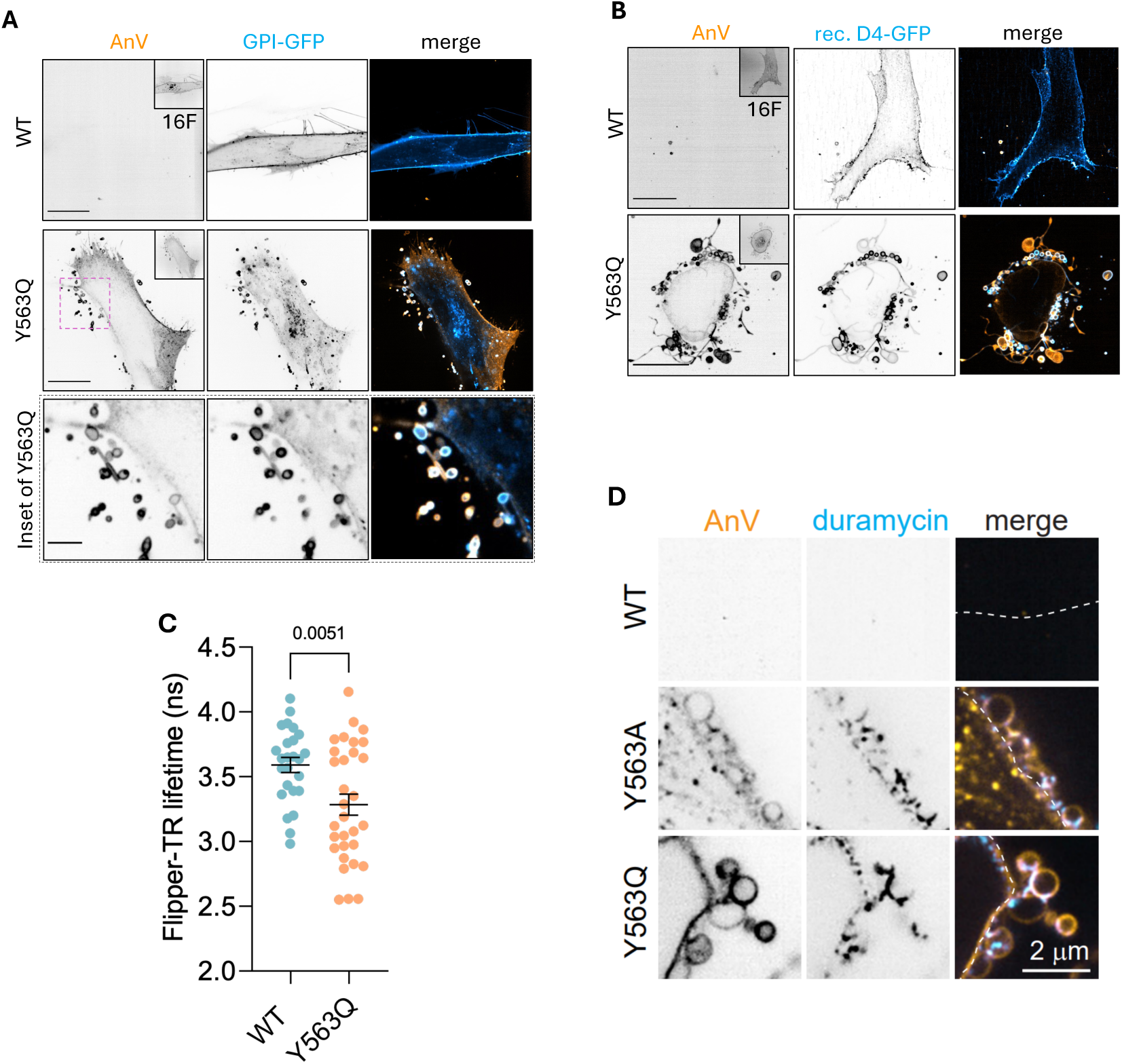
Scrambled lipid bilayers have expansive lipid-segregated domains associated with vesiculation. A) WT/YQ TMEM16F-mCherry expressed together with GPI-GFP in primary human fibroblasts and labelled with AnV. Scale bars: 20 μm; inset scale bars: 5 μm. B) WT/YQ TMEM16F-mCherry expressed in primary human fibroblasts then incubated with recombinant D4-GFP and AnV. Scale bars, 20 μm. C) Quantification of FLIM imaging of Flipper in TMEM16F WT and YQ expressing HeLa cells and quantification. D) WT/YA/YQ TMEM16F inducible HeLa cells labeled with AnV and iFluor555-duramycin. Scale bars: 2 μm

It is conceivable, however, that the excess cholesterol staining of the vesicles is a consequence of their leakiness and reflects access of the D4-GFP probe to both aspects of the bilayer. Because cholesterol redistributes rapidly across the bilayer, it may relocate preferentially to the inner monolayer if the scramblases cause net transfer of phospholipids to the outer leaflet. The inner leaflet phospholipids are thought to contain unsaturated acyl chains and their scrambling could alter lipid packing in the outer leaflet, potentially driving cholesterol inward.

The effect of the active scramblases on lipid packing was tested using Flipper, an amphiphilic probe with two planar fluorophores that undergo orientation-dependent energy transfer. The lifetime of the emission is dependent on the alignment of the two fluorophores, which is sensitive to lipid packing and/or to the lateral tension[27]. In our experimental setting such measurements need to be taken with caution since the vesicles generated by the scrambled cells are permeable, allowing Flipper access to both sides of the bilayer, confounding the interpretation. We therefore measured Flipper fluorescence lifetime only in regions of the cells that had not undergone vesiculation and hence were intact, i.e. not leaky. Comparing the lifetime of the Flipper probe in flat regions of the plasmalemma suggested that lipid packing was lower in the outer leaflet of cells expressing the active scramblase compared to WT TMEM16F (Fig. 5C). These observations are in accordance with the anticipated appearance of unsaturated phospholipids on the outer leaflet, explaining also the paucity of cholesterol from planar regions of the scrambled plasma membrane.

Like PS, phosphatidylethanolamine (PE) and PE plasmalogen are found almost exclusively in the inner leaflet of normal, healthy cells but are thought to appear on the outer surface during apoptosis. We therefore predicted that PE would also be translocated outward by the Y563A or Y563Q variants of TMEM16F. This was tested with labeled duramycin, a cyclic peptide known to bind selectively to the PE headgroup. As expected, recombinant iFluor555-labeled duramycin bound to the outer surface of cells expressing the active scramblases, while normal cells remained unstained (Fig. 5D). The insertion of the highly unsaturated PE into the outer leaflet is consistent with the reduced packing reported by Flipper. It is noteworthy, however, that the distribution of PE differed from that of PS, with the former accumulating at the base of budding vesicles yet often excluded from the vesicles themselves, while PS was generally present throughout the vesicle surface. Curiously, TMEM16F itself was also segregated away from the forming vesicles (e.g. Fig. 1B and insets to Figs 2C and 3A), suggesting that it partitions preferentially to more fluid regions of the membrane. These observations are discussed in greater detail below.

## Discussion

Vesiculation of the plasma membrane is a hallmark of most forms of cell death, notably apoptosis. Nascent ApoVs are sites of first contact with phagocytes [28] and, if released, can have impacts on the neighbouring tissue microenvironment. Despite their importance, surprisingly little was known about the events leading to apoptotic vesiculation. The experiments described above were intended to investigate the underlying molecular mechanisms. Isolating the contribution of lipid scrambling from the multiple effects of Ca^2+^ was a key objective. Using mutant forms of TMEM16F we were able to show that lipid scrambling –in the absence of [Ca^2+^] changes– suffices to generate myriad vesicles with features resembling ApoVs. Vesiculation was not accompanied by discernible changes in the cytoskeletal structure, which prompted us to search for alternative mechanisms underlying membrane curvature and scission. The acute segregation of lipids and lipid-linked proteins on the outer monolayer of scrambled cells was the salient finding: cholesterol and GPI-linked proteins accumulated markedly in the vesicles and in the convex regions of the membrane where they appeared, while PE seemed to be present in the concave neck of the forming vesicles. Precise determination of the lipid concentrations on the outer leaflet of the vesicles that had detached from the cells was hampered by the realization that they are leaky, allowing the probes used access to both aspects of the membrane. Instead, we were better able to compare lipid abundance on the outer leaflet of the main body of the cells, which remained impermeant to the probes. Clearly, cholesterol and GPI were depleted from the bulk surface of scrambled cells, compared to their abundance in untreated cells, consistent with their lateral segregation into the nascent vesicles and, in the case of cholesterol, with compensatory redistribution to the cytosolic leaflet.

We propose that the segregation of exofacial lipids underlies the generation of ApoV-like vesicles following lipid scrambling. Unlike the phospholipids and sphingomyelin present on the outer leaflet of normal cells, PS and PE are frequently endowed with kinked, unsaturated acyl chains that are thought to partition into more fluid domains of the membrane. Upon scrambling, PS and especially PE will segregate away from lipid rafts enriched in glycolipids, GPI-linked proteins, and sphingomyelin; we did not find evidence for the appearance of the latter on the inner monolayer, suggesting that it is poorly scrambled, if at all. A fraction of the highly abundant cholesterol, which can traverse the membrane rapidly and spontaneously, must relocate to the inner monolayer to fill the spaces vacated by PS and PE. Nevertheless, we believe that the remaining exofacial cholesterol partitions away from the bulk [29] regions of the membrane to associate with the rafts. The appearance of considerable amounts of phospholipids with fluid, unsaturated acyl chains must prompt the coalescence of the remaining exofacial rafts, whose large headgroups confer onto them convex (outward) curvature. The scission of these outward bulging regions is likely facilitated by the observed accumulation of PE at their base. Because of its bent (unsaturated) acyl chains and small headgroup, PE is believed to adopt a conical shape that imposes a concave (inward) curvature[30]. The lateral apposition of the convex rafts and concave PE-rich regions would foster vesicle scission. The small but significant outward hydrostatic pressure known to exist across the membrane of mammalian cells may further promote the vesiculation events, particularly if the incipient curvature leads to detachment of the convex regions from the cytoskeleton. The modest change in surface charge of the inner leaflet and/or redistribution of the phosphoinositides that anchor the skeleton to the bilayer may be involved. However, dissociation of the bilayer from the cytoskeleton is neither necessary nor sufficient to induce vesiculation. We reached this conclusion from experiments where we completely ablated the F-actin cytoskeleton with latrunculin and then acutely activated lipid scrambling with calcium ionophores. Pronounced vesiculation was observed after –but none before– addition of the ionophore (not illustrated). While not required for scramblase-induced vesicles that mimic ApoVs, it is important to reiterate that the generation of the much larger ApoBDs does involve actomyosin contractility and is hence impaired by blebbistatin, which was without effect on ApoV release (Fig. 3).

TMEM16F plays multiple roles in development and immunity. The scramblase is implicated in the formation of the placenta, where it promotes the fusion of trophoblasts in the blastocyst. TMEM16F is equally important for the fusion of osteoclast progenitors to form multinucleated osteoclasts. While the mechanisms are not entirely clear, PS that becomes exposed on the outer leaflet is likely to be recognized by receptors and/or fusogens on the surface of the opposing cell, bringing the two membranes into close proximity. The bulging of the membrane, afforded by the topology described in this report, is anticipated to aid in such a step, serving to expose membrane beyond the bulky exofacial glycocalyx.

Our findings highlight the role of TMEM16F in ApoV formation. Whether or not ApoVs can deliver important signals at a distance –including “find me” signals to attract phagocytic cells– is worthy of future investigation. It is clear, however, that the leaky ApoVs play different roles from the exosomes, secreted extracellular vesicles that contain RNA and other cytosolic molecules. The mechanisms that underlie the elevated permeability of ApoVs are therefore worth contemplating.

## Acknowledgments

We thank Dr. Greg Fairn (University of Dalhousie) for reagents and advice. J.C.M.H. is supported by the Deutsche Forschungsgemeinschaft (SFB 1557 - project P14 and project 512783018). S.A.F. is the recipient of a Canada Research Chair and supported by grants PJT-169180 from the Canadian Institutes of Health Research (CIHR) and RGPIN-2022-04485 from the Natural Sciences and Engineering Research Council of Canada (NSERC). S.G. was supported by grant FDN-143202 from CIHR. All graphics were created with BioRender.com.

## Materials and Methods

### Primary cells and cell lines

Primary fibroblasts from healthy human male and female donors were from the Coriell Institute Biobank (sample ID: GM07545 and GM23973). HEK293T (female), HeLa (female), and Jurkat (male) cells were purchased from American Type Culture Collection (ATCC).

Primary fibroblasts, HEK293T, and HeLa cells were maintained in DMEM (Wisent Inc.) supplemented with 10% FBS (called complete DMEM). Jurkat cells were maintained in RPMI 1640 (Wisent Inc.) supplemented with 10% FBS (called complete RPMI). WT control, SMS 1&2 DKO, and nSMAse 2 KO HeLa cells were described previously[21] and also maintained in complete DMEM. Cultured cells were maintained under 5% CO_2_ at 37°C.

### Generation of doxycycline-inducible mTMEM16F-expressing cell lines

WT HeLa and Jurkat cells were co-transfected or electroporated with SB100 and each of the eGFP-tagged pSBTet-Pur-mTMEM16F WT, Y563A, and Y563Q plasmids at a 1:1 ratio. HeLa cells were transfected using Fugene 6 (Promega); Jurkat cells were electroporated using a Neon system (Thermo Fisher Scientific) according to the manufacturer’s protocol. 48 h after transfection/electroporation, cells were switched to and maintained in complete medium containing 1 µg/mL puromycin. After antibiotic selection, cells were used as polyclonal populations. To induce mTMEM16F expression in the HeLa and Jurkat lines, the cells were treated with 1 μg/mL doxycycline hydrochloride (Sigma-Aldrich Cat#D9891) for the indicated times.

### DNA plasmids

eGFP-tagged WT, Y563A and Y563Q mTMEM16F plasmids were generous gifts of Dr. Huanghe Yang (Duke University)[16]. TMEM16F-mCherry and mutant forms were generated by Gibson Assembly (Cat # E2611L, New England Biolabs, Ipswich, MA, USA) of an mCherry tag, PCR-amplified from pmCherry-N1 (Cat # 632523, Takara Bio, Shiga, Japan) using primers TL01: 5’ GTGCGGCCAAAACTCGAAGCACCGGTCGCCACCATGGTGAGCAAGGGCGAGGAG and TL02: 5’ GATTATGATCTAGAGTCGCGGCCGCTCTACTTGTACAGCTCGTCCATG, as well as TMEM16F-GFP that was digested with AgeI and NotI to remove the GFP tag. eGFP-tagged pSBTet-Pur-mTMEM16F WT, Y563A, and Y563Q were created by inserting eGFP-tagged WT, Y563A and Y563Q mTMEM16F into SfiI-digested pSBTetPur vector (Addgene # 60507) by using the Gibson Assembly technique. eGFP-tagged WT, Y563A and Y563Q mTMEM16F inserts were generated by PCR amplification using TLpSBTet_F (5’-cttcctaccctcgaaaggcctctgaggccaccATGACTAGGAAGGTCCTGCTG) and TLpSBTetGFP_R (5’ gtctatcgatggaagcttggcctgacaggccTTACTTGTACAGCTCGTCCATG) primers and eGFP-tagged WT, Y563A and Y563Q mTMEM16F plasmids. The eGFP-tagged pSBTet-Pur-mTMEM16F WT, Y563A, and Y563Q plasmids were sequence-verified using the following primers:

- AS714: 5’ GTCGATTAGGCGTGTACGGTG
- m16F-aa237: 5’ GGAATCTACAAAGCAGCGTTTCC
- m16F-aa772: 5’ GCTACATCAACAACACTCTCTC
- GFPf-aa98: 5’ CCATCTTCTTCAAGGACGACG
- m16F-aa410: 5’ CGT TGA GCT ACA GCA AGA GG
- m16F-aa500: 5’ CCA CAT CCA TCACAG CCT CC

Other plasmids used in this study are pCMV T7 SB100 (Addgene plasmid # 34879), GCaMMP6s (a gift from Douglas Kim & GENIE Project; Addgene plasmid # 40753; http://n2t.net/addgene:40753; RRID:Addgene_40753), PH-PLCδ1-mCherry is from Hammond et al., 2014, R-Pre GFP (Addgene plasmid # 17274 ; http://n2t.net/addgene:17274 ; RRID:Addgene_17274) was created previously in our laboratory [19], LifeAct-EGFP was from Dyche Mullins (Addgene plasmid # 58470), and EqtSM-GFP was created previously [21], Galectin 3-mCherry was from Hemmo Meyer (Addgene plasmid #85662), GPI-GFP was created previously [31].

### Transfection

Appropriate plasmids were transfected into primary human fibroblasts and HeLa cells using FuGENE HD and FuGENE 6 (Promega), respectively. Transfection followed the manufacturer’s protocol. For transfection with FuGENE HD, transfected cells were washed and replaced with fresh medium after 5 h of transfection. Plasmids were electroporated into Jurkat cells using a Neon electroporation system (Thermo Fisher). In this case, electroporated cells were washed and replaced with fresh medium after 24 h. In all cases in which plasmids were expressed transiently, experiments were completed within 24-48 h after transfection.

### Western blotting

WT HeLa and TMEM16F (WT, Y563A, and Y563Q mutants)-inducible HeLa cells were cultured in 6-well plates for 2 days to reach 90% confluence, with two wells designated for each cell line. WT HeLa cells were treated with 2 µM staurosporine (STS) overnight. Cell lysates were collected using 4x Laemmli buffer supplemented with 2-mercaptoethanol. The lysates were then loaded into 8% SDS-PAGE gels. After the samples were separated in SDS-PAGE gels, they were transferred to PVDF membranes using the Bio-Rad wet transfer system 100 V for 2.5 h. The membrane was blocked with 5% BSA in Tris-buffered saline (TBS) for 1 h at room temperature. The membrane was then incubated overnight at 4°C with appropriate primary antibodies diluted in blocking buffer. Primary antibodies used were: 1:1000 spectrin α2 (Santa Cruz, sc-48382), 0.4 μg/mL GAPDH (Sigma; Cat# MAB374). After primary antibody incubation, the membranes were washed 4 times (5 min each) with TBS containing 0.1% Tween-20 (TBST). Subsequently, the membranes were incubated for 1 h at room temperature with appropriate secondary antibodies diluted in blocking buffer. The secondary used was 1:7000 Horseradish Peroxidase (HRP)-conjugated donkey anti-mouse antibody (Jackson ImmunoResearch). After secondary antibody incubation, the membranes were washed 4 times with TBST (10 min/wash). Chemiluminescence signals were developed using ECL^TM^ Prime Western Blotting Detection Reagents (Cytiva) and detected using a ChemiDoc XRS System (BioRad).

### Recombinant lipid probes

Recombinant D4-GFP was a generous gift from Dr. Greg Fairn[25] and was used at 1:200 dilution. Recombinant eGFP-PH-PLCδ1 was created as described previously[32]. Cells were labeled with 0.2 μM recombinant eGFP-PH-PLCδ1 for 5 min at room temperature. Cells were washed once with Annexin V-binding buffer (AnV buffer) (10 mM HEPES, 140 mM NaCl, 2.5 mM CaCl_2_, pH 7.4) before imaging. iFluor 555-conjugated duramycin was made by labeling recombinant duramycin (Santa Cruz; sc-239840) with iFluor™ 555 succinimidyl ester (AAT Bioquest; 1028(AAT)). Briefly, 44 μL of 50 mM duramycin was mixed with 7 μL of 10 mg/mL iFluor™ 555 succinimidyl ester and 5 μL of 0.7 M borate buffer, pH 8.4 for 1 h at 25°C in a shaker. The solution was then dialyzed with 2K MWCO Pierce™ Slide-A-Lyzer® Dialysis Cassettes (Thermo Fisher; 66205) for 48 h in PBS in the cold room. PBS was refreshed twice during dialysis. After dialysis, iFluor 555-conjugated duramycin was collected and stored at - 20°C. iFluor 555-conjugated duramycin was used at 1:600 dilution. Alexa Fluor 647-conjugated AnV (Invitrogen; Cat# A23204) was used at 1:400 dilution for imaging.

### Ca^2+^ imaging

Human fibroblasts were co-transfected with 8 µg of either mCherry-tagged WT, Y563A or Y563Q mTMEM16F plasmid and 5 µg of GCaMP6s plasmid using electroporation. Medium was changed after 24 h, and experiments were conducted 48 h post-electroporation. Cells were imaged in AnV buffer. GCaMP6s signal was recorded before and after 5 µM ionomycin addition. Basal Ca^2+^ levels were quantified by normalizing GCaMP6s signals before ionomycin addition to that after ionomycin addition.

### Vesicle enumeration by electronic sizing

TMEM16F (WT, Y563A, and Y563Q)-inducible HeLa and Jurkat cells were cultured to 70% confluence and then treated with 1 µg/mL doxycycline hydrochloride (Sigma-Aldrich Cat# D9891) for 24 h. For treatment with other compounds, cells were treated with either 10 µM blebbistatin, 10 µM Q-VD-OPh, or 1 µM calpeptin together with 1 µg/mL doxycycline hydrochloride (Sigma-Aldrich, Cat#D9891) also for 24 h. For experiments where the role of Ca^2+^ was assessed, cells were treated with 1 µg/mL doxycycline hydrochloride in either complete medium or Ca^2+^-free complete medium for 15 h.

Vesicles released from HeLa cells were recovered by first collecting the medium where cells had been incubated and setting this aside. Adherent cells were then detached by trypsinization. After that, the collected medium was added back to the cells to neutralize the trypsin. 0.5 mL of cell suspension of each condition was diluted with 9.5 mL Isoton II Diluent (Beckman Coulter) before analysis with a Multisizer Coulter Counter (Beckman Coulter). The total number of particles ranging from 1.5 – 60 µm was measured, and then the number of particles sized between 1 – 5 µm was normalized to the number of particles sized 8 – 60 µm (representing the total cell population).

### Flow cytometry

TMEM16F (WT, Y563A, and Y563Q)-inducible Jurkat cells were cultured to 70% confluence and then treated with 1 µg/mL doxycycline hydrochloride for 24 h. Cells were pelleted and washed once with cold HBSS. After that, the cells were labeled with 1:1000 cholera toxin B CF594 in HBSS for 30 min in 4°C. Cells were washed 3 times with cold HBSS. DAPI and 1:700 Alexa Fluor 647-conjugated AnV were added to the samples. After 15 min on ice, the samples were analysed by flow cytometry.

### Microscopy

Samples were imaged using a Quorum spinning disc mounted on a Zeiss Axiovert 200M microscope using either 63x or 25x objectives. The spinning disc microscope was coupled to an ORCA-Fusion BT CMOS camera (C9100-13, Hamamatsu). Image acquisitions were done with Volocity software (Perkin-Elmer). Imaging data were exported, analyzed, and quantified using Volocity and Fiji (NIH) software. Fluorescence signals of lipid biosensors such as R-Pre GFP and PH-PLCδ were quantified by normalizing the emission of probes on the membrane to that of the cytosol.

### Flipper-TR measurements

HeLa cells were transfected with either mCherry-tagged WT or Y563Q mTMEM16F plasmids using FuGENE HD. Medium was changed after 5 hours, and experiments were conducted 24 hrs post-transfection. Transfected cells were washed once with AnV buffer and then incubated with 1 μM Flipper-TR in AnV buffer for 15 min at room temperature. Cells were washed once with AnV buffer before imaging. The lifetime of Flipper-TR on the bulk membrane of WT or Y563Q mTMEM16F-expressing cells was measured using a STELLARIS 8 laser scanning confocal microscope equipped with a pulsed white light laser (Leica).

### Quantification and statistical analysis

Number of experiments and cells quantified are indicated in the individual Figure Legends. *n* represents individual experiments using cells with different passage numbers. Prism software (GraphPad) was used to perform all statistical analyses. An unpaired two-tailed Student’s t-test was used to compare the two groups. One-way ANOVA or two-way ANOVA with Tukey’s multiple comparisons test was used to compare multiple groups. Unless otherwise stated, error bars represent mean ± standard error of the mean (SEM). P values smaller than 0.05 were considered statistically significant.

---

**Supplemental Figure 1.**
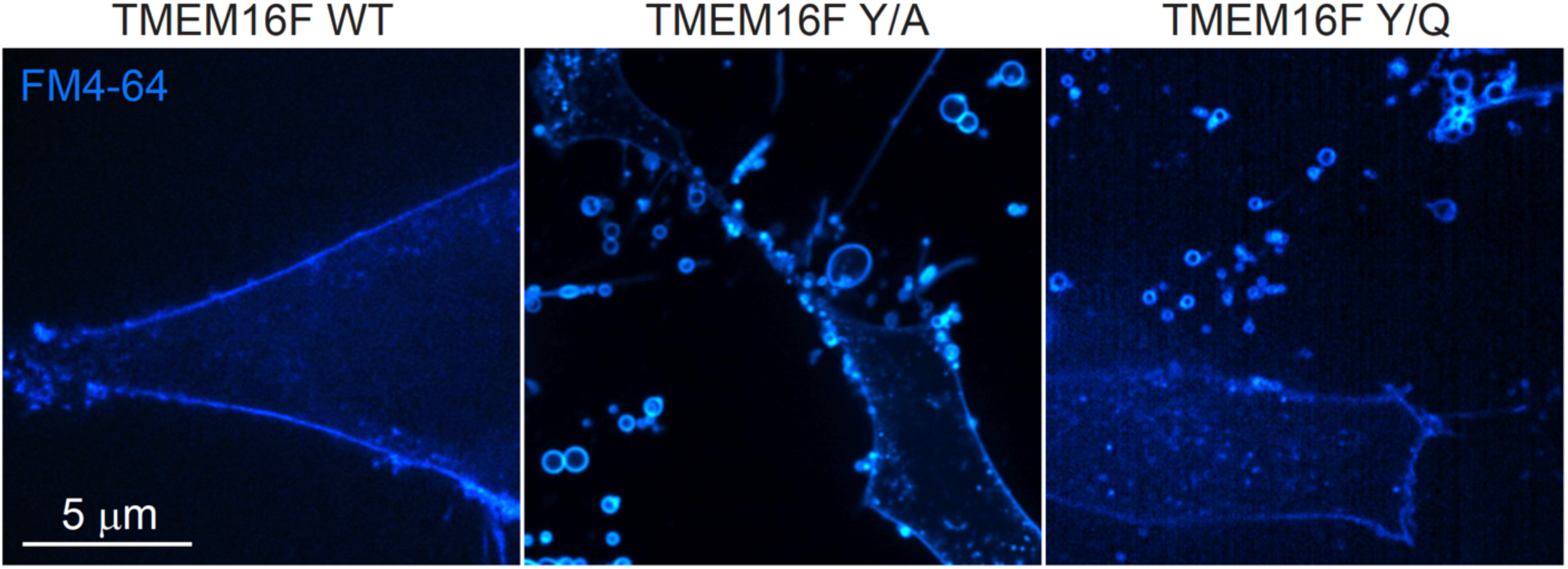
TMEM16F induced vesicles labeled with FM4-64. HeLa cells expressing indicated forms of the scramblase are labeled with FM4-64, which shows brighter labeling of the scramblase-generated vesicles, suggesting they are leaky.

